# Comparing multi- and single-sample variant calls to improve variant call sets from deep coverage whole-genome sequencing data

**DOI:** 10.1101/078642

**Authors:** Suyash S. Shringarpure, Rasika A. Mathias, Ryan D. Hernandez, Timothy D. O’Connor, Zachary A. Szpiech, Raul Torres, Francisco M. De La Vega, Carlos D. Bustamante, Kathleen C. Barnes, Margaret A. Taub, Behalf of the CAAPA consortium

## Abstract

**Motivation:** Variant calling from next-generation sequencing (NGS) data is susceptible to false positive calls due to sequencing, mapping and other errors. To better distinguish true from false positive calls, we present a method that uses genotype array data from the sequenced samples, rather than public data such as HapMap or dbSNP, to train an accurate classifier using Random Forests. We demonstrate our method on a set of variant calls obtained from 642 African-ancestry genomes from the The Consortium on Asthma among African-ancestry Populations in the Americas (CAAPA), sequenced to high depth (30X).

**Results:** We have applied our classifier to compare call sets generated with different calling methods, including both single-sample and multi-sample callers. At a False Positive Rate of 5%, our method determines true positive rates of 97.5%, 95% and 99% on variant calls obtained using Illumina’s single-sample caller CASAVA, Real Time Genomics’ multisample variant caller, and the GATK Unified Genotyper, respectively. Since most NGS sequencing data is accompanied by genotype data for the same samples, our method can be trained on each dataset to provide a more accurate computational validation of site calls compared to generic methods. Moreover, our method allows for adjustment based on allele frequency (e.g., a different set of criteria to determine quality for rare vs. common variants) and thereby provides insight into sequencing characteristics that indicate data quality for variants of different frequencies.

**Availability:** Code will be made available prior to publication on Github.

## 1 Introduction

Whole-genome sequencing has become increasingly common as a method to query genetic differences between individuals, both for population genetic studies and studies of genetic factors contributing to clinical phenotypes (Koboldt et al., 2013). Methods for translating sequenced fragments into individual genotype calls have gone through a period of active development, and many different options are available (DePristo et al., 2011; Liu et al., 2013; Cleary et al., 2014). Each of them must account for the occurrence of sequencing errors in determining whether a genetic variant is present in a particular sample, a condition that becomes especially challenging with lower sequencing depth, or in the case of a variant that has either never been seen in a given population or which is very rare.

One key decision researchers make when choosing a variant caller is whether to use a single-sample or multi-sample calling algorithm, with the argument in favor of multi-sample calling including the fact that information can be borrowed across individuals at sites of shared genetic variation. However, little work has been done to characterize the differences between the sets of variants generated by different calling algorithms, making it challenging for researchers to make a principled choice when designing an analysis pipeline. The increased computational burden of performing multi-sample calling across a large cohort means that benefits of such a calling method should be understood before carrying out this phase of a study. In addition, while genotype callers usually include some measure of quality or call confidence as part of their output, room for improvement remains in terms of better characterizing true variant calls from false positives.

In this manuscript, we present a method for characterizing variant call sets produced with different calling algorithms, in order to better select a call method suited to a particular project. We also present a method for assessing variant call quality that incorporates external genotyping array data for each subject, in order to build and train a classifier which distinguishes true variant calls from false positives. Such external array data is quite often generated either to accompany whole-genome sequencing data for sample quality control purposes, and we leverage this additional resource to improve sequencing call quality. We demonstrate our method on a set of variant calls obtained from 642 high-coverage African-ancestry genomes from the The Consortium on Asthma among African-ancestry Populations in the Americas (CAAPA), sequenced to high depth (30X).

## 2 Results

To illustrate our method, we first identified SNPs on chromosome 22 from 642 high-coverage samples of African-American ancestry from The Consortium on Asthma among African-ancestry Populations in the Americas (CAAPA) (Mathias et al., 2016) using three variant calling algorithms:

1. CASAVA from Illumina
2. Population caller from Real Time Genomics (RTG)
3. HaplotypeCaller from the Genome Analysis Toolkit (GATK)

For Illumina, variants were called on each sample individually and then merged into a combined-sample callset whereas for RTG and GATK, variants were identified jointly across all 642 samples. In all cases, the same set of 642 aligned read files (BAM files) was used. The filters used to obtain the final callsets from the raw calls are described in more detail in the Methods section.

Experimental validation of putative variants is expensive and difficult to perform for thousands of variants. Therefore, we focus on variants identified in a single sample for which we have a technical replicate, allowing variants to be partially validated.

### 2.1 Differentiating between callsets from different algorithms

Figure 1 shows a Venn diagram of the overlap between the three callsets for a single individual. From the figure, we can see that about 65.3% of all variants are (43,291 out of 66,335) are found by all three methods. Nearly 19.1% of all variants (10,385 out of 66,335) are found by only two out of three methods while 15.6% (12,658 out of 66,335) are found by only one method. To determine if any genomic features affected the ability of methods to detect a given variant, we trained a Random Forest classifier to identify which subset of methods would identify a variant with given genomic features. A detailed description of the genomic features used can be found in the Methods section and in the Supplementary Information.

**Figure 1.**
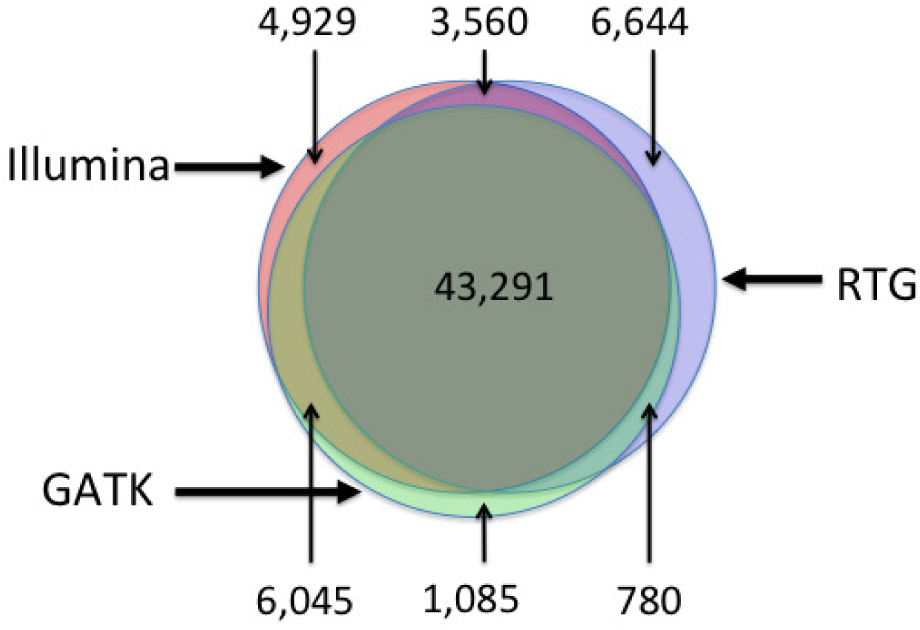
Overlap between the three callsets consisting of variants on chromosome 22 for our individual of interest.

Table 1 shows the result of the classifier on the entire variant set. The classifier overall has 1.2% error rate, suggesting that it is possible to predict with very high accuracy which methods will find a given variant given its genomic features. Figure 2 shows which genomic features are important for classifying a given variant. We can see that the frequency of the variant in the combined-sample callset is important for determining whether a variant is found in an indivdidual callset or not. Other genomic features such as the variant quality score, mapping quality, quality by depth and allele count also affect the accuracy of the classifier. The accuracy of the classifier depends not only on the coverage in the chosen individual but also on the total coverage at the given variant across all individuals. We found that using strictly genomic features without frequency information produced an error rate of 22.85%, which is lower than the error rate of a classifier that always predicts the majority class (error rate 34.7%), indicating that even without the important differentiator of how many other people in a specific call set carry a variant, there are differences in features that distinguish the call sets.

**Figure 2.**
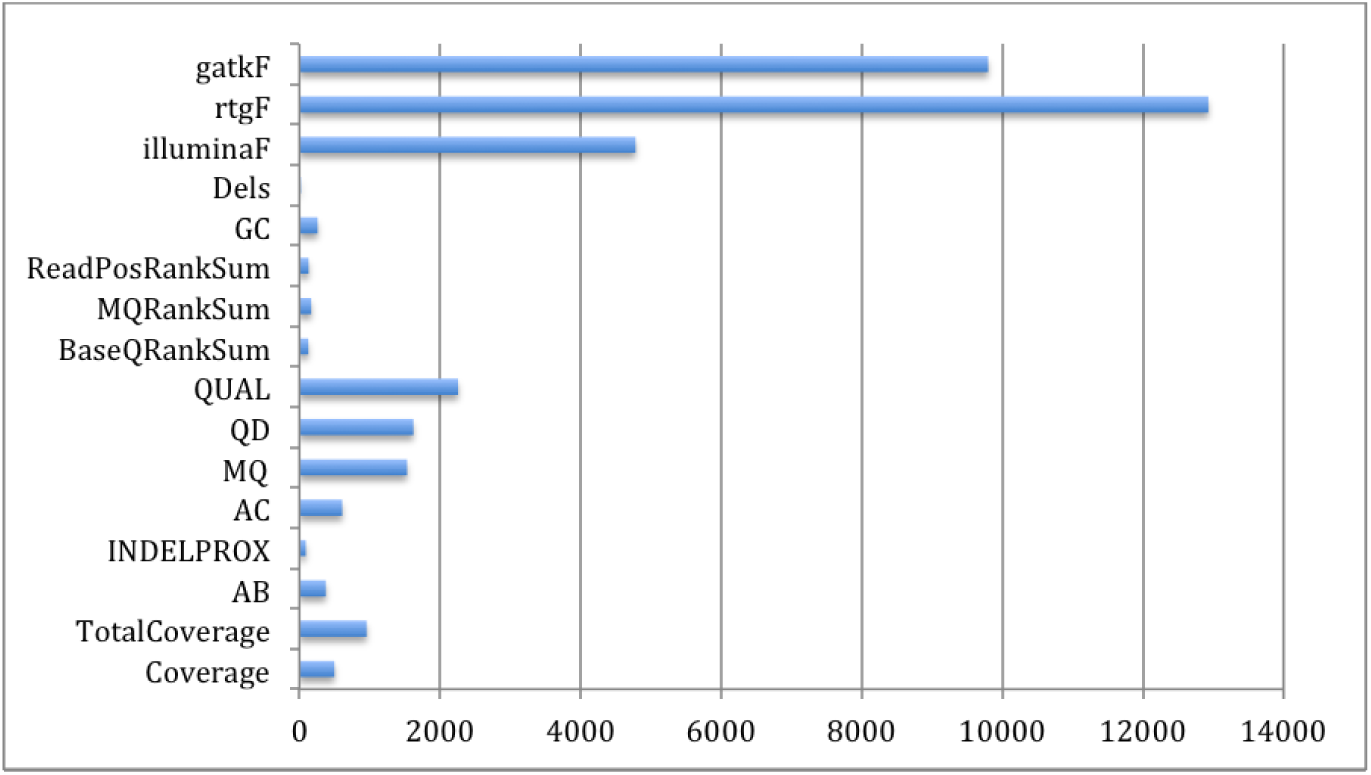
Feature importance for the Random Forest classifier distinguishing calls made by different calling algorithms. Scale on the x-axis is unitless but indicates relative importance of the different features.

**Table 1.** Confusion matrix for classifying variants. Rows show the “true” labels of variants depending on which methods find a variant. Columns show the predictions from our Random Forest classifier. The overall error rate of the classifier is 1.2%.

|  | Illumina only | Illumina + RTG | Illumina + RTG + GATK | Illumina + GATK | RTG only | RTG + GATK | GATK only | Error Rate |
| --- | --- | --- | --- | --- | --- | --- | --- | --- |
| Illumina only | 4787 | 44 | 4 | 15 | 79 | 0 | 0 | 3% |
| Illumina+RTG | 4 | 3326 | 1 | 0 | 229 | 0 | 0 | 7% |
| Illumina+RTG+GATK | 0 | 0 | 43290 | 1 | 0 | 0 | 0 | 0% |
| Illumina+GATK | 0 | 0 | 16 | 6019 | 0 | 0 | 10 | 0% |
| RTG only | 19 | 201 | 0 | 0 | 6424 | 0 | 0 | 3% |
| RTG+GATK | 0 | 0 | 95 | 0 | 4 | 681 | 0 | 13% |
| GATK only | 0 | 0 | 0 | 73 | 3 | 2 | 1007 | 7% |

### 2.2 Identifying true variant sites within a single callset

One feature of our data set is that in addition to whole-genome sequencing data, we obtained genotype microarray data from the Illumina HumanOmni2.5 BeadChip (Omni). For each of the three callsets, we used the Omni genotype data to learn a Random Forest classifier that could predict whether or not a variant site detected in a callset was truly variable or not. Using the Omni data, we declare a site to be truly variable if it is variable in both the sequencing call set and based on the Illumina array data; if the site is only variable in the sequencing data set but not based on the array data, we declare it to be a false positive. For this task, we used a larger set of features than that used earlier (see Methods). Figure 3 shows the importance of various features for the three Random Forest classifiers trained. As before, allele frequency was the most important determinant for the Illumina callset and the second most important for the GATK callset, while haplotype score, a measure of sequence complexity around the site, was the most important feature for the RTG callset. Features of just slightly less importance were allele balance (AB, highest relative importance for GATK), total coverage, haplotype score, GC content and coverage. Figure 4 shows the receiver-operating characteristic (ROC) curves for the three classifiers. We see that all three classifiers have high sensitivity and specificity (true positive rate > 0.95 with false positive rate =0.05), with area under the curve (AUC) greater than 0.99. This suggests that truly variant sites for each callset can be determined statistically using a machine-learning algorithm (similar to the Gaussian mixture model used by GATK’s variant quality score recalibration (VQSR) scoring scheme, but on an individual basis).

**Figure 3.**
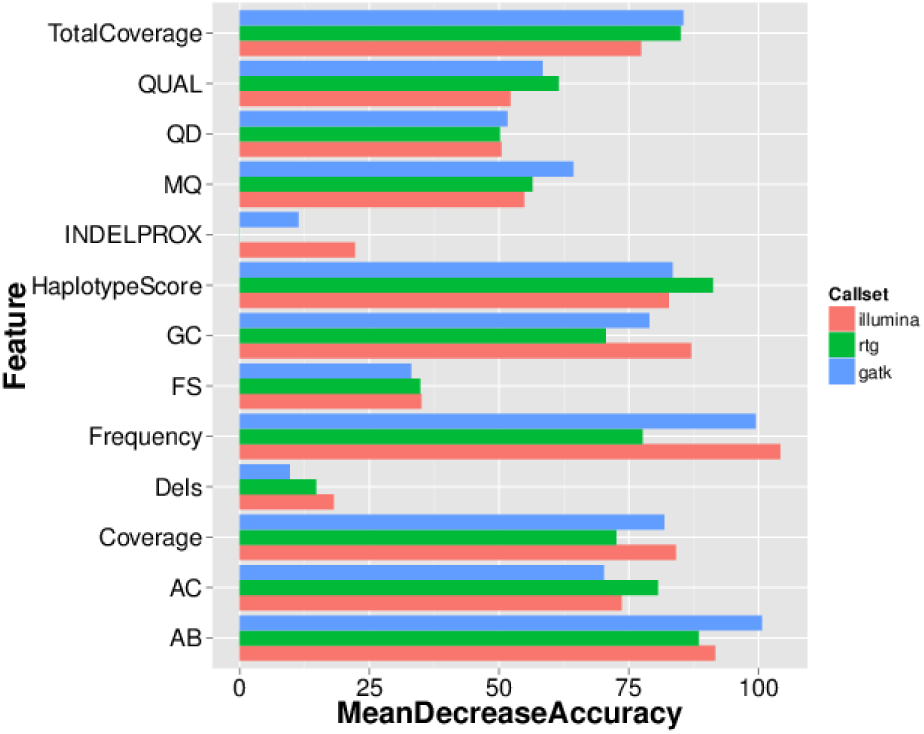
Feature importance for callset-specific classifiers based on Omni genotype data. Note that the frequency features refer to the estimates of the allele frequency from the callset being studied.

**Figure 4.**
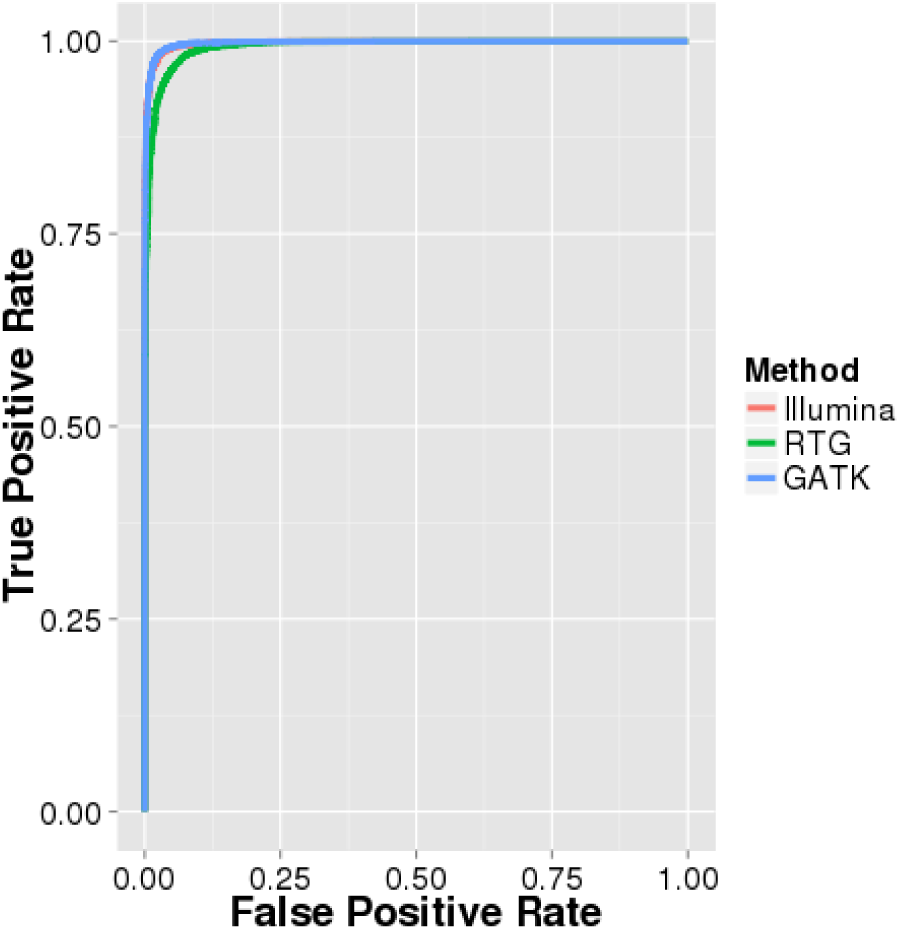
ROC curves for callset-specific classifiers based on Omni genotype data.

We evaluated the validity of our classifier scores by stratifying sites by the callsets they were found in to compare score distributions for sites with varying degrees of concordance across callsets. Figure 5 shows the results of this analysis. Considering the results for the Illumina classifier (left panel in Figure 5), we can see that sites found in all three callsets have higher classifier scores (n=43291, median score = 0.96) than sites found in two of three callsets (n=9605, median score = 0.92), with *p* < 2 × 10^−16^ for a one-sided t-test comparing these groups. Sites found only by the Illumina callset have the lowest classifier scores (n=4929, median score = 0.78), with *p* < 2 × 10^−16^ for both one-sided t-tests comparing them to the other two categories. Assuming discovery by many variant callers to be a signal that a site is truly variable, this suggests that our learned classifiers can predict truly variable sites accurately.

**Figure 5.**
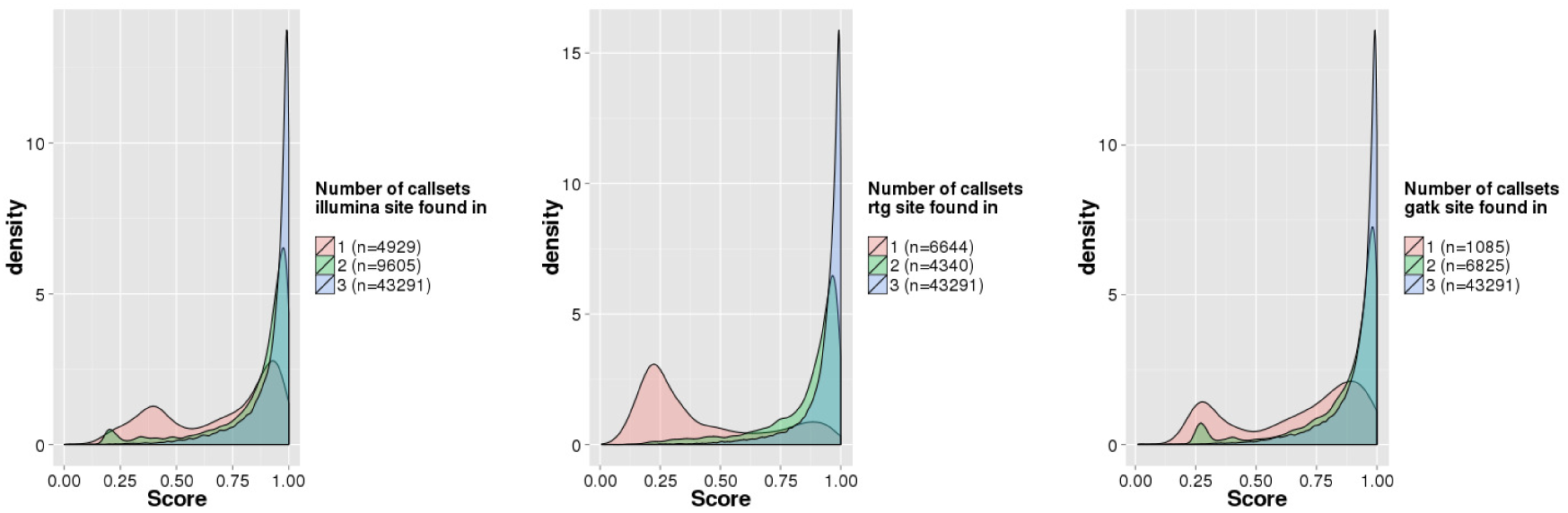
Callset-specific classifier scores for all sites, stratified by the callsets in which the site was found. Shown are calls from Illumina (left), RTG (center) and GATK (right). Colors represent the number of callsets a particular variant was observed in.

For orthogonal validation, we used a technical replicate and generated variants using the three variant callers. For each site in the original callset, we were therefore able to ascertain whether it was found in both the original and replicate callsets or just one of the two. Figure 6 shows the results of this analysis. We can see that sites appearing in both replicates have a higher classifier score than those appearing only in one replicate (for the one-sided t-test, *p* = 1.063 × 10^−5^ for illumina, *p* < 2.2 × 10^−16^ for RTG and GATK).

**Figure 6.**
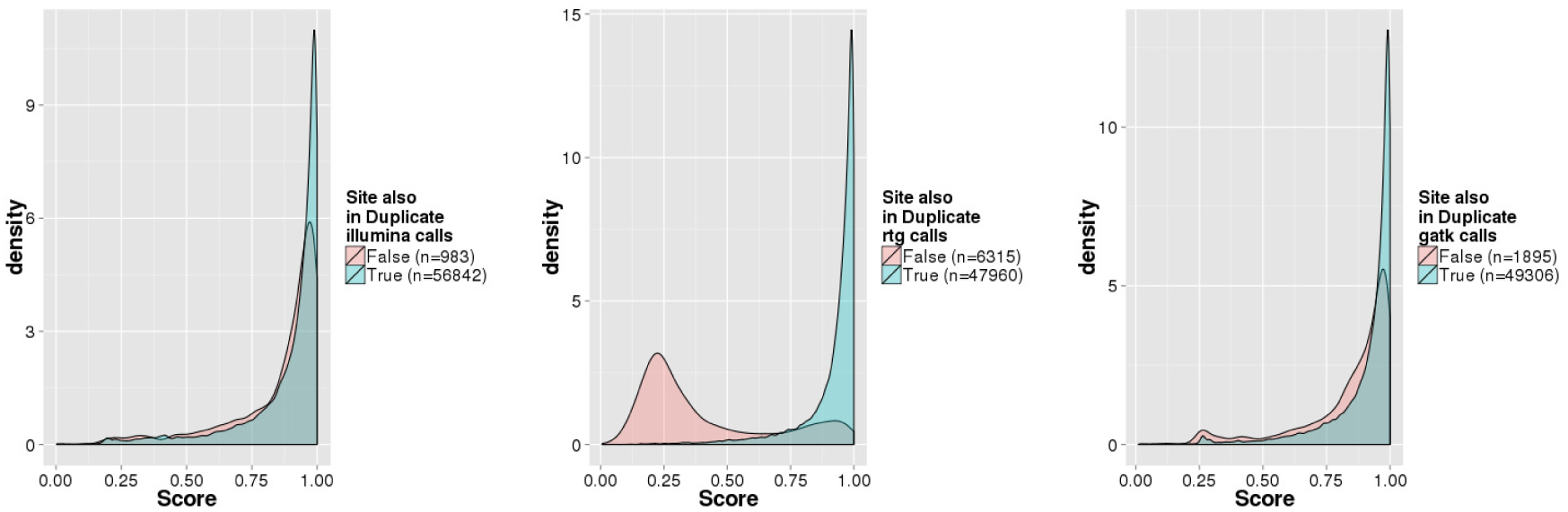
Callset-specific classifier scores for all sites stratified by whether the site was found in both replicates or only one replicate. Shown are calls from Illumina (left), RTG (center) and GATK (right). Colors represent the number of replicates a particular variant was observed in.

For each callset, we used the fitted callset-specific classifer to predict whether all discovered sites within the callset were variable. Table 2 shows the results of this prediction.

**Table 2.**
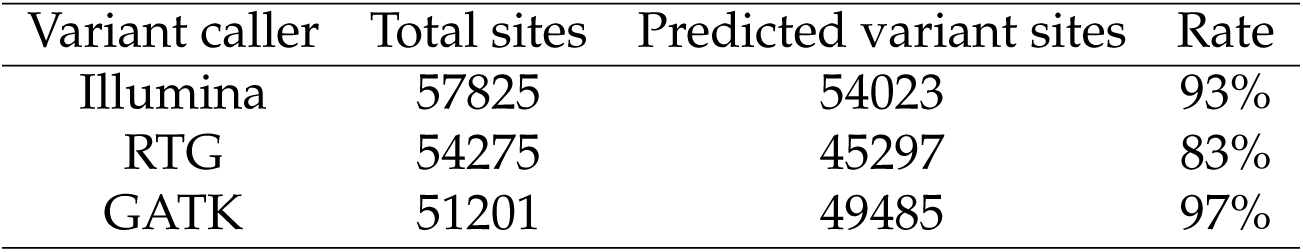
Predicted number of true variant sites from three different callsets using fitted callset-specific classifiers.

| Variant caller | Total sites | Predicted variant sites | Rate |
| --- | --- | --- | --- |
| Illumina | 57825 | 54023 | 93% |
| RTG | 54275 | 45297 | 83% |
| GATK | 51201 | 49485 | 97% |

We see that of the three methods, the Illumina callset has the largest number of sites predicted to be variable. The GATK callset has the highest proportion of sites predicted to be variable, which is expected since the callset was filtered using VQSR before our analysis. Overall, all three callsets have a high proportion of variants predicted to be true calls by our classifier.

## 3 Methods

For our analyses, we used alignment files (BAMs) that we received from Illumina after sequencing for the 642 samples. Variant calls were obtained from the alignment files using three different pipelines as follows:

- Illumina: Raw calls were filtered first by removing any variants near centromeres or other high copy regions by filtering based on depth of coverage, and second by removing variants falling into regions of segmental duplication. In addition, individual genotype calls were set to missing which had call quality scores below 20 or depth of coverage below 7. The merged multi-sample VCF was then filtered for SNPs that failed Hardy-Weinberg equilibrium (HWE, *p* < 10^−6^). Calling was performed at Illumina, where the sequencing was performed.
- RTG: We used the population caller from RTG (version 3.2) to jointly obtain variant calls for the 642 samples. The multisample VCF was filtered using the Adapative Variant Rescoring scheme (minAVR = 0.025). The resulting VCF was then filtered for SNPs that failed HWE with *p* < 10^−6^.
- GATK: We used the GATK UnifiedGenotyper (version 2.7.4) to jointly obtain variant calls for the 642 samples. The multisample VCF was filtered using VQSR (sensitivity=94%). The resulting VCF was then filtered for SNPs that failed HWE with *p* < 10^−6^.

Illumina also provided us with genotype microarray data from the Illumina HumanOmni2.5 BeadChip (Omni). There are 33234 SNPs on the Omni array for chromosome 22. Of those, 4123 SNPs (12%) have *MAF* < 0.01, 10221 SNPs (31%) have *MAF* < 0.05 and 23012 (69%) have *MAF* ≥ 0.05.

### 3.1 Distinguishing between callsets

To distinguish between the three callsets, we used a random forest classifier implemented in the “randomForest” package (Liaw and Wiener, 2002) written in R (R Core Team, 2014). Random forest classifiers are collections of decision trees that allow non-linear interactions between features and are robust to over-fitting (Breiman, 2001). We used 1000 trees for our analysis due to the large number of variants.

#### 3.1.1 Features selected

We used a number of genomic features for the callset-specific classifiers. Many of them were chosen to be among the features used in GATK’s VQSR scoring scheme. Supplementary Table 1 describes the features used.

### 3.2 Call-set specific classifiers

#### 3.2.1 Features selected

We used a number of genomic features for the callset-specific classifiers. Supplementary Table 2 describes the features used. Based on results from the previous analysis, read allele imbalance measured using a RankSum test (such as BaseQualityRankSum) were not very informative for classification and were replaced by a Fisher test score of allele balance for this analysis.

#### 3.2.2 Ground truth data

For our classification task, we used the Omni genotype calls as ground truth. Therefore, we were able to use the sites overlapping between the Omni SNPs and the variant calling output from the sequencing data as our labeled set. Sites that were heterozygous or homozygous alternate in the Omni genotype calls for the chosen individual were lableled ‘1’ while sites which were homozygous reference were lableled ‘0’.

#### 3.2.3 Unbalanced class problem

On intersecting the sequencing variant calls with Omni genotype data to obtain labeled sites, we observed that the ratio of the number of sites lableled ‘1’ to those labeled ‘0’ was nearly 100:1 (11506:184 for illumina, 10376:170 for RTG and 11465:184 for GATK). In classification tasks, this can be problematic since it can bias the classifier towards increasing accuracy by always predicting the majority class.

To avoid this problem we used the SMOTE method (synthetic minority over-sampling technique) by Chawla et al. (2002) implemented in the R software (Torgo, 2010). This method oversamples the minority class by creating synthetic examples of the minority class from existing examples. It also undersamples the majority class for improved performance.

## 4 Discussion

Here, we present a method of using random forests to both characterize different variant call sets and to assess call quality taking into account a wide variety of data features in a flexible way. We show that for sequencing data like ours, e.g., with relatively deep (30x) coverage, both single-sample and multi-sample calling methods provide calls with very good accuracy. We illustrate our method on a single sample for which we had a technical replicate to provide further insight into the behavior of our classifier.

To further explore differences between the callsets, we also looked at results stratified by allele frequency bin. Here there were some differences in performance, with different thresholds chosen for classifying rare and common sites resulting in nearly 800 additional sites being predicted as variant (Supplementary Table 3). In light of these results, we conclude that the optimal calling method to apply may depend on what the intended use of the variant calls is, with different applications (e.g., population level vs variant specific analyses) best served by different calling methods. For example, recent work (Han, Sinsheimer, and Novembre, 2014) discusses potential biases in site-frequency spectrum estimation that can result from low-coverage sequencing data where rare variants are more likely to be missed. For an application like this, multi-sample calling would be most appropriate to leverage information from many samples. However, in general applications or when sequencing coverage is high as it is in our data set, we have not observed a large impact on downstream results comparing the different call sets. As part of the analysis in the CAAPA flagship publication, we compared some downstream results generated with both the Illumina single-sample callset and the RTG multi-sample callset and found no difference in the overall patterns seen in the count of deleterious alleles by individual or by group (see Supplementary Information of (Mathias et al., 2016)). We do note that our false positive rate of 5% may be considered high for applications to disease genetics, and suggest that the standard practice of validating any interesting findings either through replication or through further genotyping should still be used. Finally, modifications to the method presented here to target a particular allele frequency class, such as modifying the quality threshold for different frequency bins, or potentially retraining the classifier on different subsets of the data split by frequency bin, are also possible if variants in a particular frequency range are of special interest in a particular application.

While collecting genotype array data is common practice to accompany whole-genome sequencing data generation for the purpose of sample verification, our work indicates that it is also valuable to have array data as an orthogonal validation data set for QC purposes and to assess overall call quality of the variant calls from the sequencing data.

As an alternative to the SMOTE method, weighting the observations in the different classes to give a more balanced training set is a possibility. We also implemented this and found that performance was not as good as what we present here (data not shown).

A potential extension to the work presented here would be to develop a method of producing a consensus call set based on the results of the classifier output for each set of variant calls. We could produce a combined result from the different calling methods using a weighted combination (e.g., weighted by the quality score assigned by our classifier) or a latent variable model. This would be similar to work presented in (DePristo et al., 2011) but with a more flexible approach to assigning weights to the various possible calls.

## 5 Conclusion

We demonstrate that differences between callsets generated by different variant callers can be explored and interpreted using machine learning methods like Random Forests. In addition, we show that Random Forests can be used to identify which variant calls from a specific callset are likely to be true.

## Acknowledgments

Funding for this study was provided by National Institutes of Health (NIH) R01HL104608.

